# SOORENA: Self-lOOp containing or autoREgulatory Nodes in biological network Analysis

**DOI:** 10.1101/2025.11.03.685842

**Authors:** Hala Arar, Jehad Aldahdooh, Payman Nickchi, Mohieddin Jafari

## Abstract

Autoregulatory mechanisms, in which proteins regulate their own activity or expression, are fundamental to biological networks but are challenging to identify systematically from literature. To address this gap, we present SOORENA (https://soorena.it.helsinki.fi/soorena/), a two-stage transformer model that predicts and classifies protein autoregulation in PubMed abstracts. SOORENA was trained on 1,332 experimentally validated abstracts and achieved 96.0 percent accuracy and 97.8 percent precision in stage one, with stage two achieving 95.5 percent accuracy and 96.2 percent macro-F1 across seven mechanistic classes. Applied to 3.34 million abstracts, SOORENA identified 85,145 publications containing autoregulatory mechanisms, yielding 97,657 protein-specific records. Integration with curated databases generated 100,065 comprehensive entries accessible via an interactive Shiny application. By systematically cataloging self-regulatory interactions, which often act as bottlenecks in dynamic network modeling, SOORENA provides a resource that supports mechanistic interpretation, model reduction, and predictive systems-level analyses. These results demonstrate that domain-specific language models can scale the discovery and curation of biologically essential self-regulatory mechanisms, bridging literature mining and systems biology.

## 1. Introduction

Autoregulation refers to a biological process in which a protein modulates its own abundance or activity through direct molecular feedback, enabling precise control of cellular function [1]. These self-regulatory processes are widespread across signaling, transcriptional, and metabolic networks, where they contribute to homeostasis, robustness, and rapid adaptation to changing environments [2]. Autoregulatory motifs are among the most common topological features observed in biological regulatory systems, particularly in transcription factor networks and enzyme signaling pathways [3–5]. Although they are often ignored in static network modeling due to limitations in current methods and algorithms, such as those used for centrality or module detection analyses, these motifs cannot be disregarded in dynamic network modeling. Notably, when represented as self loops in network models, they cannot be eliminated by any known model reduction techniques [6,7]. This underscores their fundamental importance in preserving the integrity and functionality of biological models.

Autoregulation operates through diverse molecular mechanisms. Enzymatic self-modification includes autophosphorylation by kinases, autoubiquitination by E3 ligases, autocatalytic reactions in metabolic enzymes, and autolysis in proteases [8–11]. At the gene expression level, transcription factors can activate or repress their own transcription, forming positive or negative feedback loops that modulate response dynamics and reduce expression noise [12–14]. These mechanisms play essential roles in human disease. Dysregulated autoregulatory signaling contributes to oncogenic kinase activation, neurodegenerative protein dysfunction, and bacterial quorum-sensing (QS) pathways that drive antimicrobial resistance, making autoregulatory proteins promising therapeutic targets [15,16].

Despite their importance, autoregulatory mechanisms remain under-characterized in public databases and are difficult to extract computationally from scientific literature. Mechanistic descriptions are often embedded implicitly in text, using heterogeneous phrasing rather than standardized vocabulary. For example, a statement such as “the kinase phosphorylates itself” implies autophosphorylation without naming the mechanism explicitly. Our keyword presence analysis supports this challenge. For example, fewer than half of autophosphorylation publications include the literal term in their abstracts, and similar gaps were observed across other mechanisms. As a result, conventional keyword-based search and rule-based text mining struggle to retrieve relevant literature when mechanistic descriptions are phrased implicitly.

Although manually curated resources such as UniProt provide high-quality mechanistic annotations [17], expert review cannot keep pace with the rapid expansion of biomedical literature. Publishing output now exceeds 1.5 million new scientific articles annually [18], meaning that reviewing even a targeted subset of 250,000 abstracts would require a tremendous amount of effort. This creates a substantial bottleneck in keeping biological knowledge current and comprehensive.

Natural language processing (NLP) approaches using neural language models have transformed information extraction in biomedicine by enabling semantic interpretation beyond surface keywords [19, 20]. Domain-specific transformer architectures (for example, PubMedBERT) capture biomedical terminology and contextual meaning more effectively than general-domain models. This facilitates the detection of specialized mechanistic relationships in text, such as autoregulation.

Here, we present Self-lOOp containing or autoREgulatory Nodes in biological network Analysis (SOORENA), a transformer-based model with two stages for automated identification and mechanistic classification of protein autoregulation in scientific literature. In the first stage, the model identifies abstracts describing any form of autoregulation. In the second stage, positive abstracts are classified into seven mechanistic categories: autophosphorylation, autoubiquitination, autocatalytic activity, autoinhibition, autolysis, autoinducer production, and autoregulation of gene expression. This architecture improves computational efficiency by restricting the fine-grained classification to relevant publications while preserving high precision in mechanism screening.

SOORENA was fine-tuned on 1,332 manually curated and experimentally validated mechanism annotations mapped from UniProt to PubMed abstracts. Because our dataset was highly unbalanced (with 711 examples of autophosphorylation but only 38 of autoinducer production), we used a weighted loss function and evaluation metrics that better captured performance on the underrepresented classes [24, 40]. These design choices enabled robust recognition of both common and rare autoregulatory mechanisms.

Finally, we deployed SOORENA across 3,340,955 PubMed abstracts, identifying 85,145 publications (2.5%) containing autoregulatory mechanisms, which yielded 97,657 protein-specific records after gene annotation extraction. Integration with 1,332 UniProt annotations and 1,076 interactions from pathway databases (SIGNOR [36], TRRUST [37], OmniPath [35]) yielded a total of 100,065 curated autoregulatory records, creating the largest resource to date for protein self-regulation in the literature. An interactive web application allows researchers to explore predictions, filter by metadata, and access ontology-linked standardized terms. SOORENA provides a practical framework to accelerate the discovery and curation of protein self-regulatory mechanisms by integrating expert-curated training data, domain-specific language models, and scalable inference tools.

## 2. Methods

### 2.1 Data Collection

We constructed a large-scale biomedical text corpus by integrating publication metadata from PubMed with manually curated annotations of autoregulatory mechanisms from UniProt. Specifically, we utilized the Swiss-Prot subset of UniProt, which contains reviewed protein entries with experimentally validated mechanism annotations [17]. Although UniProt references a vast number of publications (estimated at over 200 million, many of which remain unverified), our dataset focuses on a high-confidence subset derived from curated entries [17]. Abstracts were retrieved via the PubMed API and used for language modeling, while UniProt served as the source of validated autoregulatory labels, which are embedded within individual protein records [39].

#### 2.1.1 PubMed Corpus

We obtained a subset of 262,819 publications from PubMed, each containing a PubMed Identifier (PMID), Title, Abstract, Journal, and Authors. These publications were randomly selected from over two million abstracts corresponding to journals indexed in UniProt to ensure representative coverage of literature relevant to protein function. The dataset included 8,607 records lacking abstracts but contained no duplicate PMIDs.

#### 2.1.2 UniProt Annotations

We extracted experimentally validated autoregulatory mechanism annotations from the Universal Protein Resource [17]. These annotations are generated by expert curators based on experimental evidence reported in the referenced publications. The UniProt dataset consisted of 1,323,976 rows and 13 metadata fields describing protein-level experimental findings. Each entry is tied to a protein via a UniProt accession number (AC) and may reference supporting literature using PMIDs embedded in the RX field. Each unique AC may have more than one reference available. Up to three fields (RP, RT, and RC, which we later named as Term_in_RP, Term_in_RT, Term_in_RC) contain mechanistic terms indicating evidence that a protein regulates its own activity or abundance. These term columns were aggregated into a single “Terms” column for model training. Of all entries, 1,823 were labeled with autoregulatory mechanisms (number of rows with a known autoregulatory mechanism), while 1,322,153 were unlabeled. A total of 181,779 rows had missing PMIDs, including 10 labeled entries, which were removed because the model required PubMed-linked text for training. Additionally, 89,620 PMIDs appeared multiple times, since each protein record occupied its own row; these were later aggregated so that each PMID corresponded to a single, combined entry. After aggregation, the dataset was reduced from 1,142,197 rows to 268,619 unique PMIDs, of which 1,374 were labeled with autoregulatory mechanisms. The reduction in labeled entries occurred because multiple protein records often referenced the same PMID, and these were merged into a single aggregated entry.

#### 2.1.3 External Resources Integration

To augment coverage of experimentally validated autoregulatory interactions, we integrated curated data from three established pathway databases: OmniPath, SIGNOR, and TRRUST. OmniPath aggregates protein-protein interactions and regulatory relationships from over 100 literature-curated sources [35]. We extracted all self-directed regulatory interactions where source and target protein identifiers matched. SIGNOR provides manually annotated causal interactions in human signaling pathways [36]. We queried SIGNOR for instances where regulator and target proteins were identical. TRRUST contains transcription factor-target regulatory relationships derived from text mining and manual curation [37]. We extracted cases where a transcription factor regulates its own gene expression. Duplicate interactions across sources were deduplicated while retaining source attribution. Integration of external databases contributed 1,076 additional validated autoregulatory cases (SIGNOR: 995, TRRUST: 61, OmniPath: 20), representing 1.1% of the final dataset. These external resources were not used for model training but were integrated into the final database for extensive coverage. External databases use varied vocabulary for equivalent concepts (e.g., SIGNOR’s “self-phosphorylation” maps to autophosphorylation, “self-cleavage” maps to autolysis). SOORENA harmonizes source-specific terms to enable unified cross-database queries.

#### 2.1.4 Gene Annotation Extraction

To associate model predictions with specific proteins, we extracted gene and protein annotations from publications using PubTator3 [38], a state-of-the-art named entity recognition system for biomedical literature. PubTator3 processes full-text articles (when available) rather than abstracts alone, providing more comprehensive entity coverage. For each publication predicted as containing autoregulatory content, we retrieved all gene/protein mentions and created individual database records for each protein-publication pair. This approach accounts for papers describing multiple autoregulatory proteins, resulting in more records than unique publications. Extracted annotations include UniProt accession numbers, gene names, protein names, and organism identifiers when available.

### 2.2 Label Construction and Data Integration

#### 2.2.1 Merging PubMed and UniProt

UniProt annotations were mapped to PubMed metadata using PMIDs as a common identifier. A join between two tables was applied to ensure that all PubMed publications were retained, even if no corresponding UniProt record existed. Mechanism labels from UniProt were added only when a PMID match was found. The resulting merged dataset contained 262,819 PubMed publications in total, of which 1,349 were labeled with autoregulatory mechanisms and 261,470 were unlabeled. The slight reduction in labeled entries from 1,374 to 1,349 occurred because a small number of labeled PMIDs from UniProt were not present in the PubMed corpus. A total of 8,607 entries from the PubMed dataset were dropped because they lacked abstracts. Only two of these were labeled, and abstracts were required for model training, leaving 1,347 labeled entries at this stage.

#### 2.2.2 Term Analysis

To characterize the labeled data, all mechanistic terms associated with autoregulatory behavior were extracted from the UniProt annotations. We identified a total of 15 unique terms across the labeled subset, which shows substantial class imbalance. The most frequent mechanism was autophosphorylation (711 instances), followed by autocatalytic activity (133), autoubiquitination (121), autoregulation (119), and autoinhibition (84). Less common terms included autolysis (41) and autoinducer production (31), while several variants, such as autokinase, autocatalysis, and autophosphatase, appeared only rarely. The ratio between the most and least frequent terms (711:1) highlights the strong skewness toward phosphorylation-related mechanisms. This is not surprising, as this is a common trend in molecular biology due to extensive kinase research. These distributions informed later normalization and filtering steps to ensure that rare but biologically meaningful mechanisms were retained during modeling.

To assess how often mechanistic terms appeared explicitly in the literature, we examined whether each annotated term occurred literally within its corresponding abstract text. The results showed wide variability across mechanism types. Terms such as autoactivation, autoinduction, and autolysis appeared in nearly all labeled publications, while others were frequently absent from the text despite being experimentally confirmed in UniProt - most notably autoubiquitination (14.0%), autophosphorylation (47.3%), and autocatalytic (41.4%). This finding highlights that many descriptions of autoregulatory mechanisms are phrased implicitly (for example, “the kinase phosphorylates itself”) rather than through standardized terminology. As a result, keyword searches or rule-based text mining miss a large portion of the relevant literature. This highlights the need for a context-aware language model that can understand implicit mechanistic descriptions.

A small subset of publications contained more than one autoregulatory mechanism within the same abstract. Specifically, 31 publications (approximately 2.3% of all labeled data) referenced multiple mechanisms, with an average of 2.1 mechanisms per publication and a maximum of 3.

#### 2.2.3 Feature Engineering

To prepare the text for model training, the title and abstract of each publication were concatenated into a single text field to provide richer contextual information for mechanism detection. The resulting field was stored as text and served as the primary input for all subsequent modeling stages. The text was then cleaned using a series of preprocessing steps: HTML entities such as “&” were decoded, URLs and email addresses were removed, and excess whitespace was normalized. These operations ensured that the model processed standardized and noise-free text while preserving important biomedical terminology.

To evaluate data quality, text length was measured as the number of characters per abstract. Only two publications contained fewer than 100 characters, indicating minimal issues with incomplete text. Both of these publications were unlabeled. Across the dataset, the mean text length was 1,348 characters (SD = 432, median = 1,351), suggesting a consistent abstract structure across publications - with no apparent outliers in length. Labeled publications were slightly longer (mean = 1,403, SD = 382; median = 1,402) than unlabeled ones (mean = 1,348; SD = 433, median = 1,351).

We further standardized the labeled data by normalizing spelling and variant forms of mechanistic terms to ensure consistent representation. Terms such as autoregulatory, autoinhibitory, autocatalysis, and autoinduction were replaced with their canonical forms (autoregulation, autoinhibition, autocatalytic, and autoinducer). After normalization, 11 unique mechanisms remained. To ensure balanced training, we retained only terms with at least 35 examples, resulting in seven mechanism classes summarized in the total facet of **Fig. 1**. Despite this filtering, a 711:38 imbalance persisted between the most and least common mechanisms.

**Figure 1.**
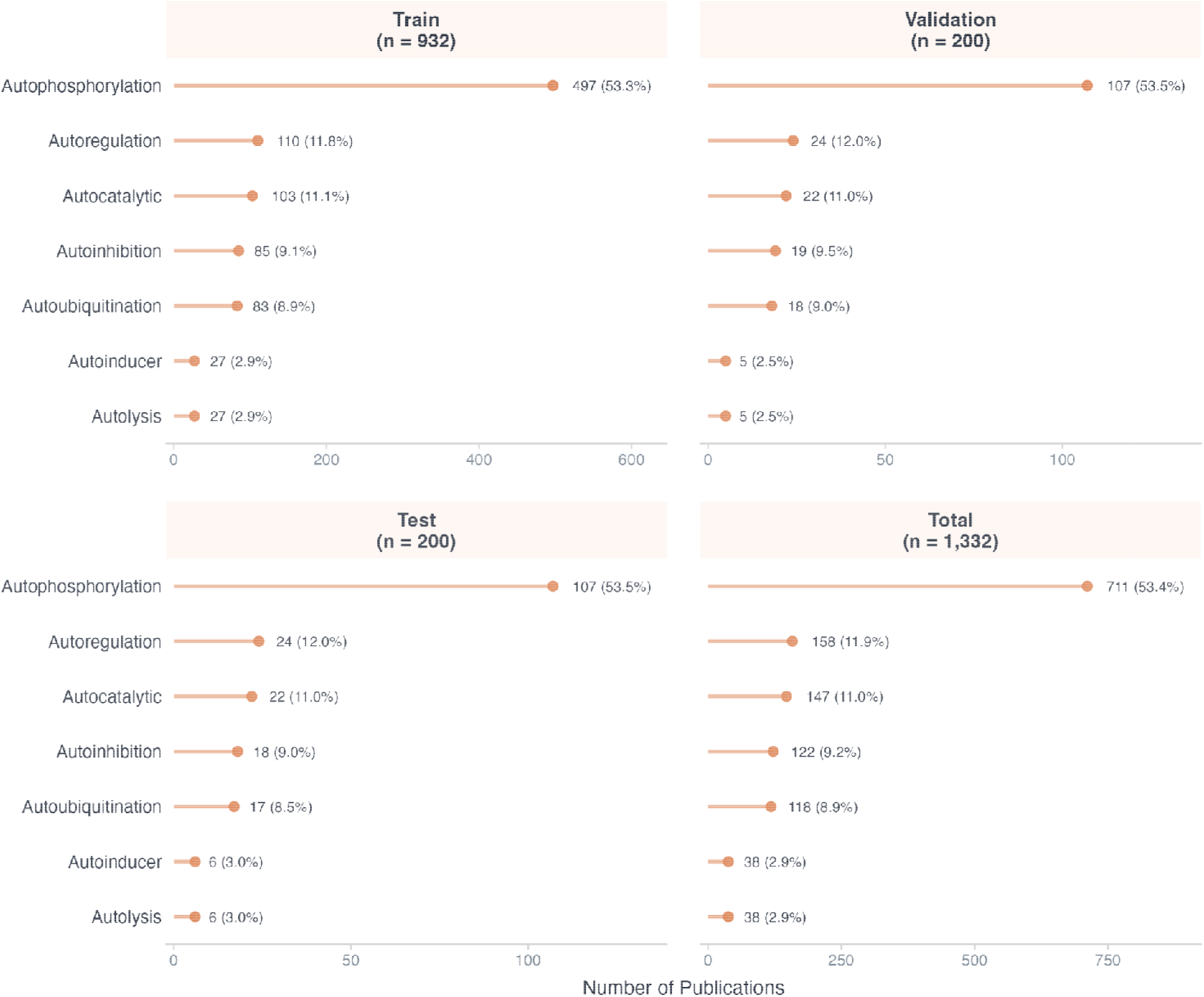
Per-class distribution of autoregulatory mechanisms across training, validation, test, and total subsets of the labeled dataset (n = 1,332). X-axes are scaled independently per panel.

A small subset of publications (18 publications, 1.3 % of labeled data) referenced multiple mechanisms, with up to three mechanisms in a single abstract. However, this subset was insufficient to train a reliable multi-label classifier because of the sample size. Therefore, in the current version of SOORENA, each publication was assigned a single primary mechanism label (the first listed term) to maintain class balance and ensure stable model training. After all cleaning, normalization, and filtering steps, the final modeling dataset consisted of 254,197 total publications, of which 1,332 were labeled with one of the seven supported mechanisms.

### 2.3 Train-Validation-Test Splits

Following dataset preparation, a total of 254,197 publications were retained, of which 1,332 contained experimentally validated autoregulatory mechanisms. These labeled examples were stratified by mechanism type to preserve class proportions across training, validation, and test sets. This stratification ensured that each mechanistic category was proportionally represented during model development, thereby minimizing sampling bias toward the mechanism class with the majority proportion (autophosphorylation).

Using this stratified procedure, 70% of labeled publications (n = 932) were assigned to the training set, while the remaining 30% (n = 400) were split evenly between validation and test sets (n = 200 each). This split was performed only on the labeled subset, meaning that the remaining 252,865 unlabeled publications were excluded from these initial splits and later incorporated during binary classification (Stage 1). The overall composition of datasets across the original stratified split, Stage 1 (binary detection), and Stage 2 (multiclass classification) is summarized in **Table 1**.

**Table 1.**
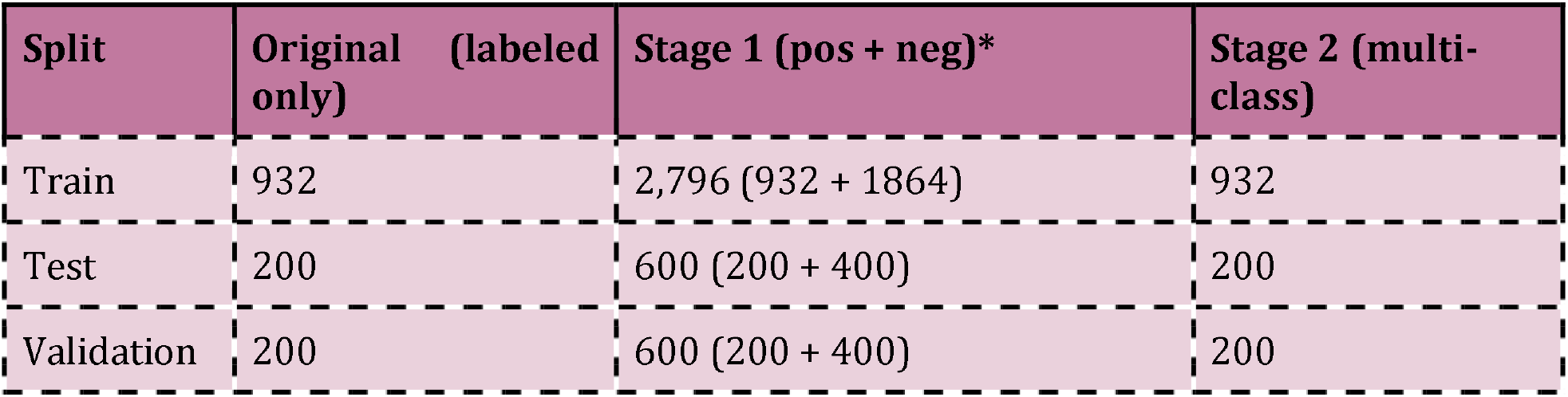
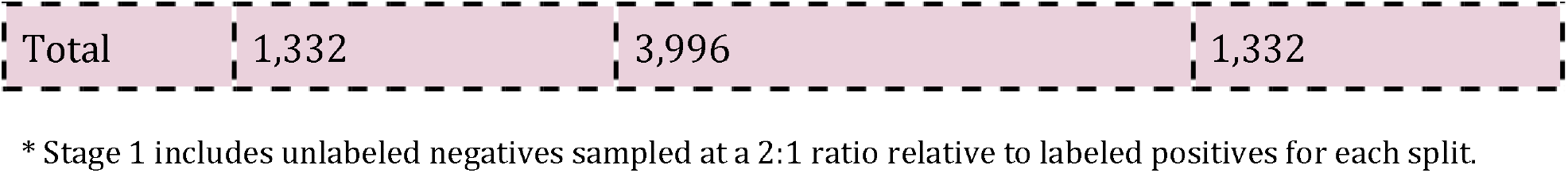
Dataset composition across original, Stage 1, and Stage 2 splits.

| Split | Original (labeled only) | Stage 1 (pos + neg)* | Stage 2 (multi-class) |
| --- | --- | --- | --- |
| Train | 932 | 2,796 (932 + 1864) | 932 |
| Test | 200 | 600 (200 + 400) | 200 |
| Validation | 200 | 600 (200 + 400) | 200 |
| Total | 1,332 | 3,996 | 1,332 |

\* Stage 1 includes unlabeled negatives sampled at a 2:1 ratio relative to labeled positives for each split.

In the original split, the data were used to train a single-stage supervised model limited to the labeled corpus. For Stage 1, the task was reformulated as binary classification, where publications describing autoregulatory mechanisms were labeled 1, and randomly sampled unlabeled publications were treated as 0. Each split was expanded by adding unlabeled negatives at a 2:1 ratio, tripling the training volume while preserving the same class balance across validation and test sets. This strategy improved the model’s ability to distinguish between mechanistic and non-mechanistic abstracts, reducing the likelihood of false positives before passing candidates to Stage 2. Because unlabeled publications were treated as negatives, a small number of true positives may exist within the negative set. However, the expected contamination rate is low: only 1,332 of 254,197 publications (0.5%) carried validated labels, and the prevalence of autoregulatory content in the broader literature is estimated at 2.5% based on deployed predictions. This level of label noise is consistent with prior work showing that BERT-based classifiers are robust to low rates of mislabeled negatives in positive-unlabeled settings [41].

For Stage 2, only the original labeled subsets (932 train, 200 validation, 200 test) were used. This stage addressed multi-class mechanism classification, predicting one of seven mechanism types. Because the dataset remained highly imbalanced (dominated by autophosphorylation), class weights were applied during loss computation to equalize contributions from minority classes. The class-wise counts within the labeled corpus are shown in **Fig. 1**.

These counts confirm that stratified sampling preserved each mechanism’s clas proportions across splits. The training subset captured ∼70% of each class, while validation and test sets maintained balanced representation for evaluation. Together, this framework produced a robust two-stage learning design: Stage 1 for binary detection of autoregulatory mechanisms across both labeled and unlabeled literature, and Stage 2 for fine-grained classification of mechanism type within the labeled subset.

### 2.4 Model Architecture

SOORENA is a two-stage trained transformer-based classification model built on PubMedBERT (microsoft/BiomedNLP-PubMedBERT-base-uncased-abstract-fulltext), a BERT-base architecture pretrained solely on PubMed abstracts and full-text articles to capture biomedical terminology and contextual semantics [20,21]. Stage 1 detects whether a publication describes any autoregulatory mechanism, serving as a screening layer to filter the broader biomedical literature. Stage 2 then assigns Stage 1-positive publications to one of seven distinct mechanism categories. PubMedBERT has demonstrated superior performance over general-domain BERT variants in biomedical entity recognition and relation extraction tasks [20].

Both stages use the same architecture: a PubMedBERT encoder followed by a dropout layer (p = 0.1) and a linear classification head. The stages are fine-tuned independently, allowing hyperparameter optimization and checkpoint selection specific to the prediction task (**Fig. 2**).

**Figure 2.**
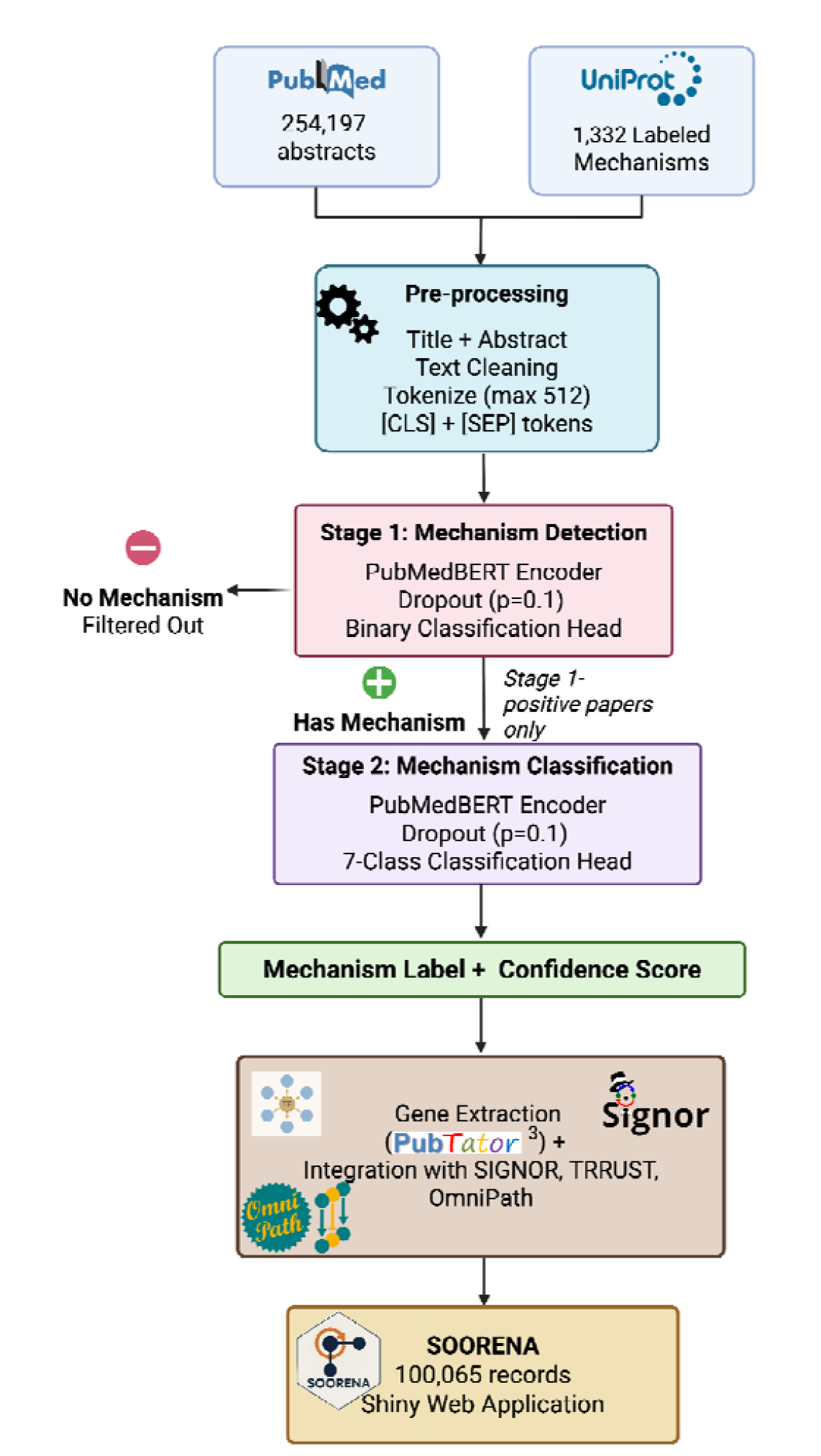
Overview of the SOORENA two-stage autoregulatory mechanism detection architecture. Created on BioRender.com.

### 2.5 Training Procedure

Both stages were fine-tuned using PyTorch 2.0 and the Hugging Face Transformers library (v4.30). Inputs were tokenized to a maximum length of 512 tokens and processed in mini-batches of 16 samples. The AdamW optimizer (learning rate of 2 × 10^−5^) was used with a linear warmup schedule applied to the first 10% of training steps. All data splits and model initialization used a fixed random seed (42) for reproducibility. All experiments were conducted on an Apple M1 Max (32 GB unified memory) without GPU acceleration. Because training was performed on CPU, results are fully deterministic and do not depend on non-deterministic CUDA operations. AdamW was chosen for its improved stability and regularization when fine-tuning transformers compared to standard Adam [22].

Stage 1 training employed binary cross-entropy loss and ran for three epochs (525 update steps). Model selection was based on the validation F1 score, which balances precision and recall to minimize false positives entering Stage 2. Stage 2 training used weighted cross-entropy loss to handle class imbalance across mechanism types and ran for four epochs (236 update steps). The class weights, inversely proportional to class frequency, are shown in **Fig. 3C**. The final Stage 2 model checkpoint was chosen based on the highest validation macro-F1 score, ensuring balanced performance across both common and rare mechanisms.

**Figure 3.**
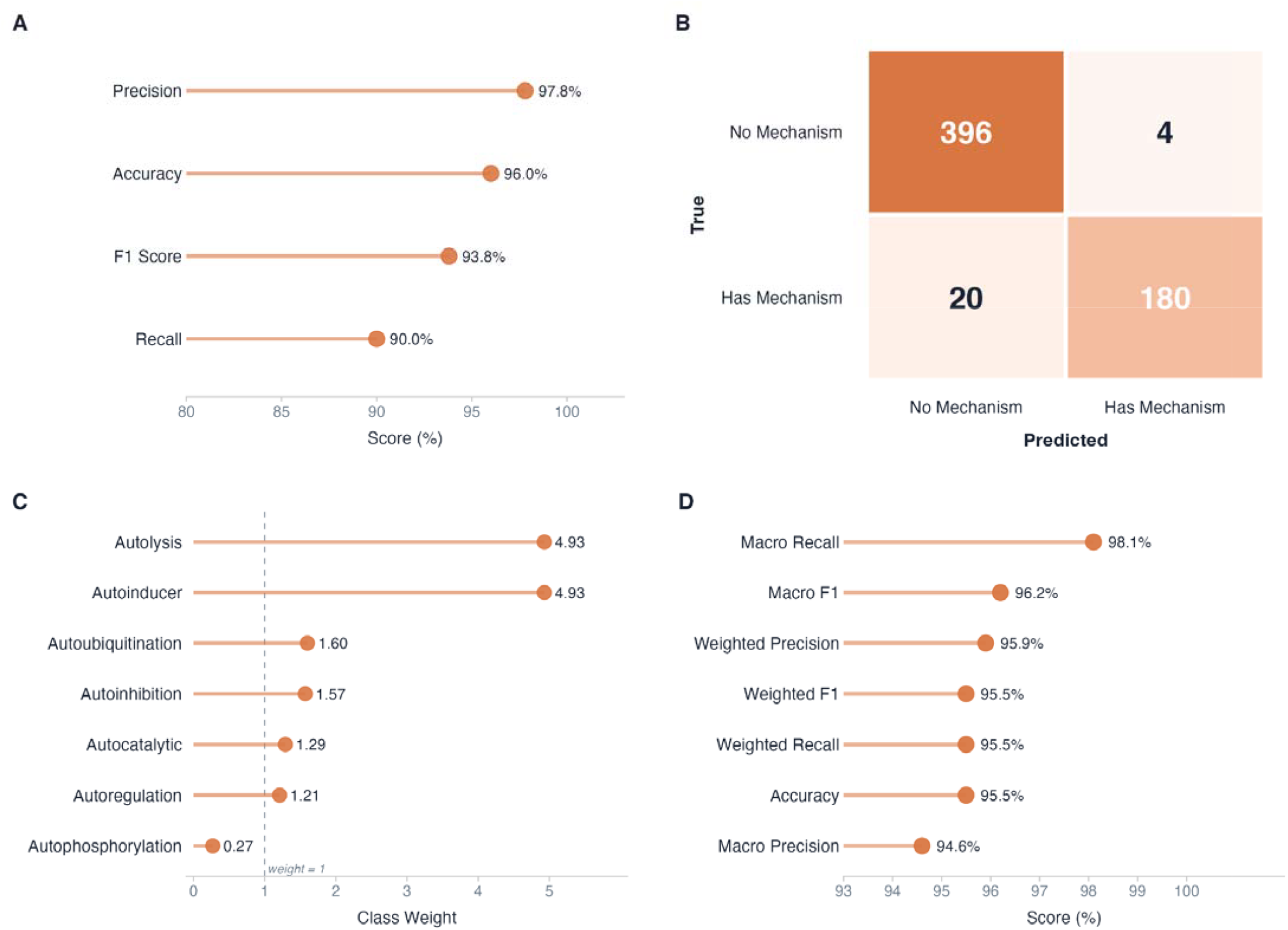
Stage 1 and Stage 2 model evaluation. (A) Stage 1 binary classifier test-set performance (n = 600), showing accuracy, precision, recall, and F1 score. (B Confusion matrix for Stage 1 predictions on the test set. (C) Inverse-frequency class weights assigned to each autoregulatory mechanism type for Stage 2 training; the dashed line indicates a neutral weight of 1.0. (D) Stage 2 multiclass classifier test-set performance (n = 200), showing accuracy and macro- and weighted-averaged precision, recall, and F1 score.

### 2.6 Evaluation

#### 2.6.1 Performance Metrics

We assessed model performance using multiple complementary metrics: Accuracy, Precision, Recall, F1 Score, Macro-F1, and Weighted-F1. For Stage 2, macro-F1 was prioritized as the primary metric because it prevents strong performance on common classes (autophosphorylation) from masking poor performance on rare ones (autolysis, autoinducer). This is critical in biomedical classification, where underrepresented mechanisms may be scientifically important despite low frequency.

#### 2.6.2 Error and Confidence Analysis

Confusion matrices were generated to inspect class-specific error patterns. Prediction confidence was used to identify uncertain classifications, especially in Stage 1, where the negative class contains label noise.

### 2.7 Computational Resources

All training and inference were performed on an Apple M1 Max (32 GB unified memory) without dedicated GPU acceleration, demonstrating that the pipeline is accessible on consumer hardware. Training time was approximately 33 minutes per epoch for Stage 1 and 12 minutes per epoch for Stage 2.

### 2.8 Ontology Development

To enable consistent terminology across heterogeneous data sources and support biologically interpretable outputs, we developed a curated ontology for the seven autoregulatory mechanism classes used by SOORENA. The ontology organizes mechanisms into a two-branch hierarchy rooted at “Autoregulatory Mechanisms” : (i) enzymatic self-modification, encompassing post-translational mechanisms (autophosphorylation, autoubiquitination, autocatalytic activity, autolysis), and (ii) gene expression regulation, encompassing transcriptional and signaling feedback (autoregulation of gene expression, autoinhibition, autoinducer production).

Each mechanism entry includes a formal definition, controlled synonyms and antonyms, related terms, and a set of core ontology relations following OWL-inspired conventions: is-a (class membership), part-of (biological process membership), regulates (functional effect), has-input and has-output (molecular participants), and occurs-in (biological context).

Each mechanism was additionally assigned a polarity indicator based on its functional direction: positive (+) for self-amplifying mechanisms (autophosphorylation, autocatalytic activity, autoinducer production), negative (–) for self-limiting mechanisms (autoinhibition, autoubiquitination, autolysis), and context-dependent (±) for autoregulation of gene expression, which can act as either a positive or negative feedback depending on the specific regulatory interaction.

The ontology also serves as a cross-database harmonization layer, mapping source-specific terminology from integrated databases to SOORENA’s canonical terms (for example, SIGNOR’s “self-phosphorylation” maps to autophosphorylation, and “self-cleavage” maps to autolysis). In the web application, each predicted record is linked to its ontology entry, providing users with standardized definitions and mechanism context alongside model predictions.

## 3. Results

### 3.1 Stage 1: Binary Classification Performance

Stage 1 classified publications as either containing or not containing evidence of an autoregulatory mechanism. On the held-out test set (n = 600), the model achieved strong performance, accurately identifying publications that should progress to mechanistic classification. Overall accuracy was 96.0%, with a precision of 97.8% and a recall of 90.0% for the positive class, yielding an F1 score of 93.8% (**Fig. 3**). The model’s high precision ensured that very few false positives were propagated into Stage 2.

Errors were dominated by false negatives, with 20 mechanism-containing publications missed and 4 publications incorrectly flagged as positive. This conservative behavior aligns with the design objective of minimizing uncertain cases passed to Stage 2. The confusion matrix (**Fig. 3**) illustrates that misclassifications did not arise from systematic bias toward specific negative subsets but rather from ambiguous descriptions in the original text.

Validation F1 improved from 90.2% at epoch-1 to a peak of 91.8% at epoch-2, followed by a slight decrease at epoch-3, indicating minor overfitting. The best epoch-2 checkpoint was therefore selected for final evaluation. Collectively, these results demonstrate that Stage 1 acts effectively as a high-precision filtering stage for biomedical corpora for fine-grained mechanistic classification.

### 3.2 Stage 2: Multi-Class Mechanism Classification

Stage 2 assigned each Stage 1-positive publication to one of seven autoregulatory mechanism categories. On the held-out test set (n = 200), the model achieved 95.5% accuracy, 96.2% macro-F1, and 95.5% weighted-F1 (**Fig. 3**). These results demonstrate robust generalization despite substantial class imbalance in the training data.

Performance across individual mechanism types was similarly strong (**Fig. 4**). Autophosphorylation, the most common mechanism, achieved 99.0% precision and 92.5% recall (F1 = 95.6%), indicating effective representation learning even when class prevalence might bias toward overprediction. Mechanisms with intermediate representation, including autoregulation, autocatalytic activity, autoinhibition, and autoubiquitination, achieved F1 scores above 91%.

**Figure 4.**
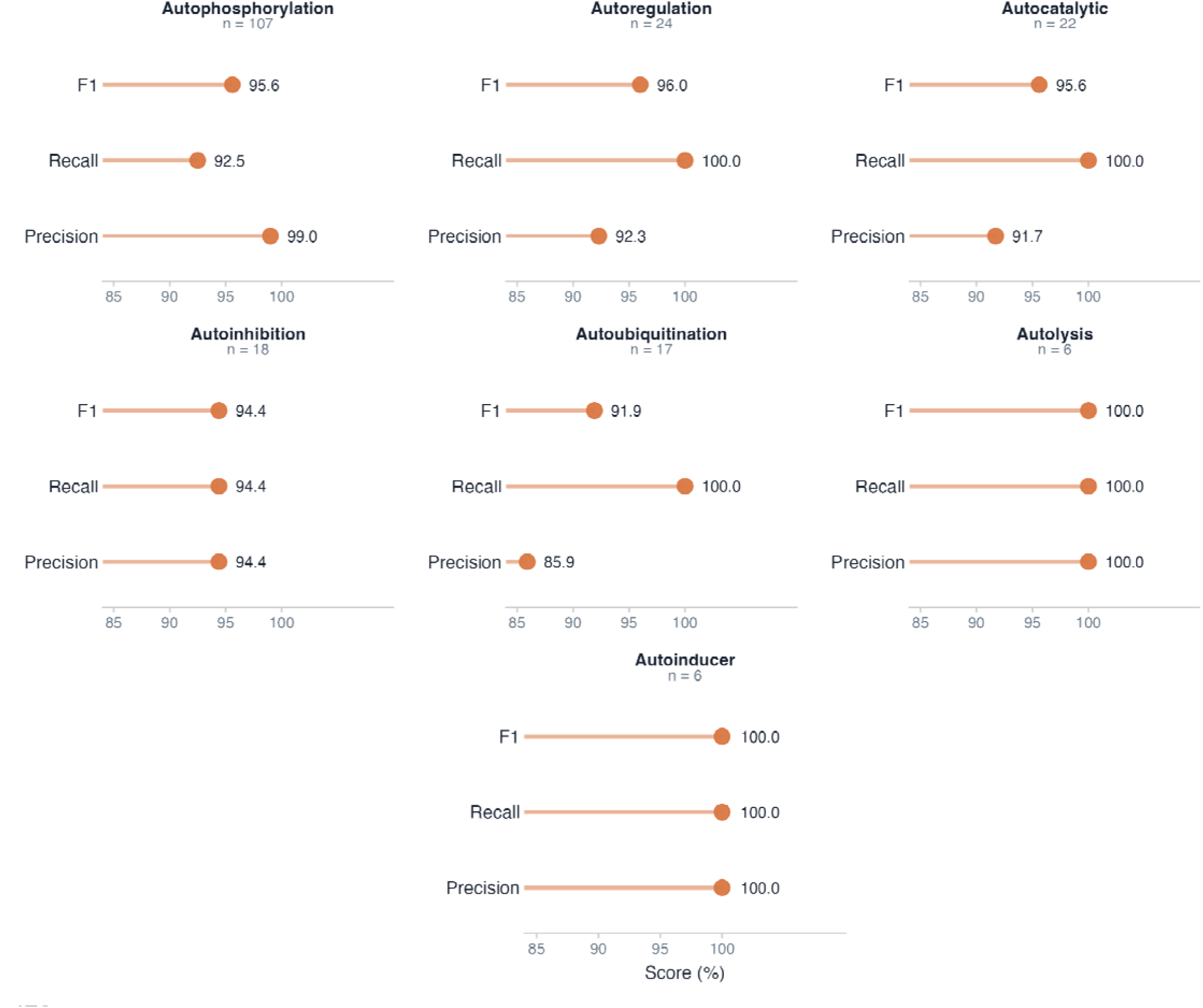
Per-class classification performance of the Stage 2 mechanism classifier on the test set. Each panel shows Precision, Recall, and F1 score for one autoregulatory mechanism class; support (n) indicates the number of test-set instances per class. The x-axis is held constant across all panels to allow direct comparison. Scores are derived from the held-out test set (n = 200 instances total).

Strikingly, the rarest mechanisms, autolysis and autoinducer production (n = 6 each), were classified perfectly (100% precision, recall, and F1). However, the small support for these classes means that a single misclassification would reduce F1 to approximately 67%. These results should therefore be interpreted with caution until validated on larger held-out samples. Nevertheless, the perfect classification of these rare classes highlights the effectiveness of weighted loss in preserving minority-class performance and preventing collapse toward majority categories. Together, these findings show that Stage 2 reliably captures mechanistic diversity across the full spectrum of autoregulatory behaviors.

### 3.3 Error & Uncertainty Analysis

Across the Stage 2 test set, the model misclassified 9 of 200 publications (4.5% error rate). Most errors occurred between mechanistically related categories, particularly autophosphorylation and autocatalytic activity, which share overlapping biochemical terminology describing self-activation mediated by the enzyme. Importantly, no misclassifications occurred between mechanistically distant categories (e.g., autolysis vs. autoinducer production), indicating that the model captures biologically meaningful distinctions rather than relying on superficial keyword patterns (**Fig. 5**).

**Figure 5.**
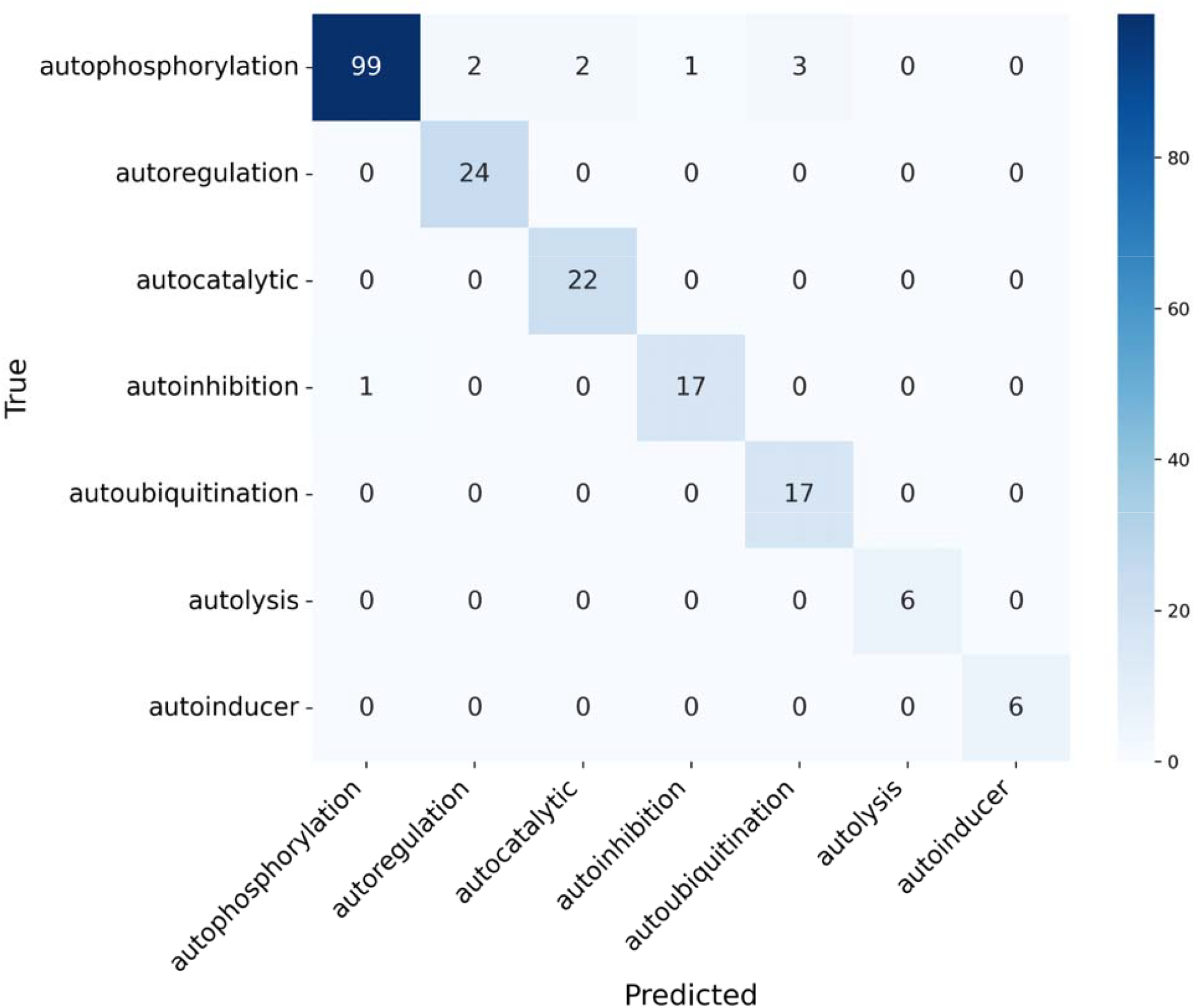
Confusion matrix for Stage 2 multi-class classification showing true versus predicted mechanism labels for 200 test abstracts.

Prediction confidence scores, representing the model’s estimated probability for each predicted class, provided additional insight into model reliability. Correctly classified Stage 2 examples had a mean confidence of 92.7%, whereas misclassified examples showed substantially lower confidence (mean: 67.3%). This separation suggests that confidence estimates can function as a practical uncertainty measure for downstream expert review.

Similar trends were observed in Stage 1. False negatives were more frequent than false positives, reflecting the model’s intentional bias toward high precision. Negative predictions also exhibited a higher mean confidence (93.3%) than positive predictions (89.2%). Together, these findings indicate that SOORENA not only achieves strong classification accuracy but also provides interpretable confidence signals that support high-confidence curation while safely flagging ambiguous literature for manual evaluation.

To assess potential cross-regulation false positives in deployed predictions, we applied pattern matching to identify cases where the abstract text explicitly describes heteroregulation (e.g., “Protein A regulates Protein B”). Of the 97,657 predicted autoregulatory publications, 3,034 (3.1%) contained explicit cross-regulation patterns. Manual inspection confirmed that these represent a source of false positives arising from a known system limitation: the model predicts whether an abstract describes autoregulation but cannot identify which protein is autoregulatory. Gene annotations are extracted separately via PubTator3 from full-text articles, and the system pairs predictions with all extracted gene mentions. Consequently, when an abstract describes heteroregulation (Protein A ⍰ Protein B), the prediction may be incorrectly associated with the target protein (B) rather than the actor (A). This pairing error accounts for an estimated 3% of deployed predictions and highlights the need for subject-protein identification in future model iterations.

### 3.4 Interactive Database: SOORENA Web Application

We developed SOORENA, an R Shiny application that integrates model predictions with experimentally validated records from multiple curated databases (**Fig. 6**). The database contains 100,065 autoregulatory interaction records: 97,657 predicted from 3,340,955 PubMed abstracts, 1,332 UniProt annotations, 995 SIGNOR interactions, 61 TRRUST interactions, and 20 OmniPath interactions. Gene annotations for predicted records were extracted using PubTator3 [38] from full-text articles. Each record is assigned a unique identifier (SOORENA-{SourceCode}-{PMID}-{Counter}), where SourceCode indicates the data source (P = Predicted, U = UniProt, S = SIGNOR, T = TRRUST, O = OmniPath), PMID is the PubMed identifier of the source publication, and Counter is a sequential integer that disambiguates multiple records sharing the same PMID and source (arising when a single publication describes more than one autoregulatory protein), enabling traceability to original publications and source databases. The application supports search, filtering, inspection, and export, and includes an embedded ontology to standardize terminology and aid interpretation.

**Figure 6.**
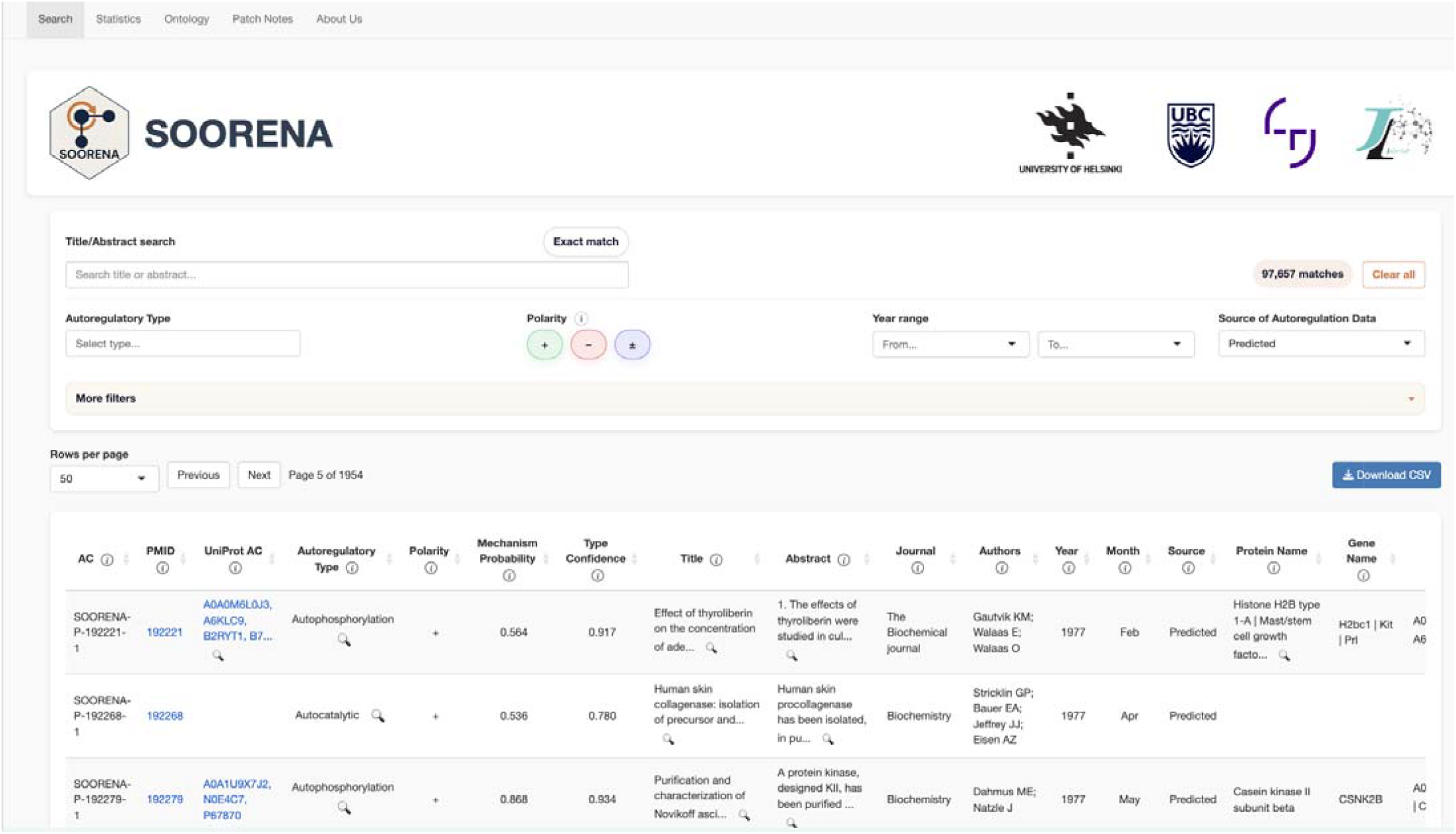
SOORENA web interface showing the Search tab, which supports field-specific filtering (e.g., Protein ID, PMID, organism, and data source), and displays interactive, sortable table columns, and CSV export functionality.

The model processed 3,340,955 PubMed abstracts and identified 85,145 publications (2.5%) as containing autoregulatory mechanisms. After PubTator3-based gene extraction, these predictions yielded 97,657 protein-specific autoregulatory records, reflecting that multiple genes may be extracted from a single publication. Integration with external databases (1,332 UniProt annotations + 1,076 interactions from SIGNOR/TRRUST/OmniPath) brought the total database to 100,065 records. The distribution of predicted mechanism types is summarized in **Fig. 7**. The table shows a dominance of autophosphorylation followed by autoubiquitination and autocatalytic activity.

**Figure 7.**
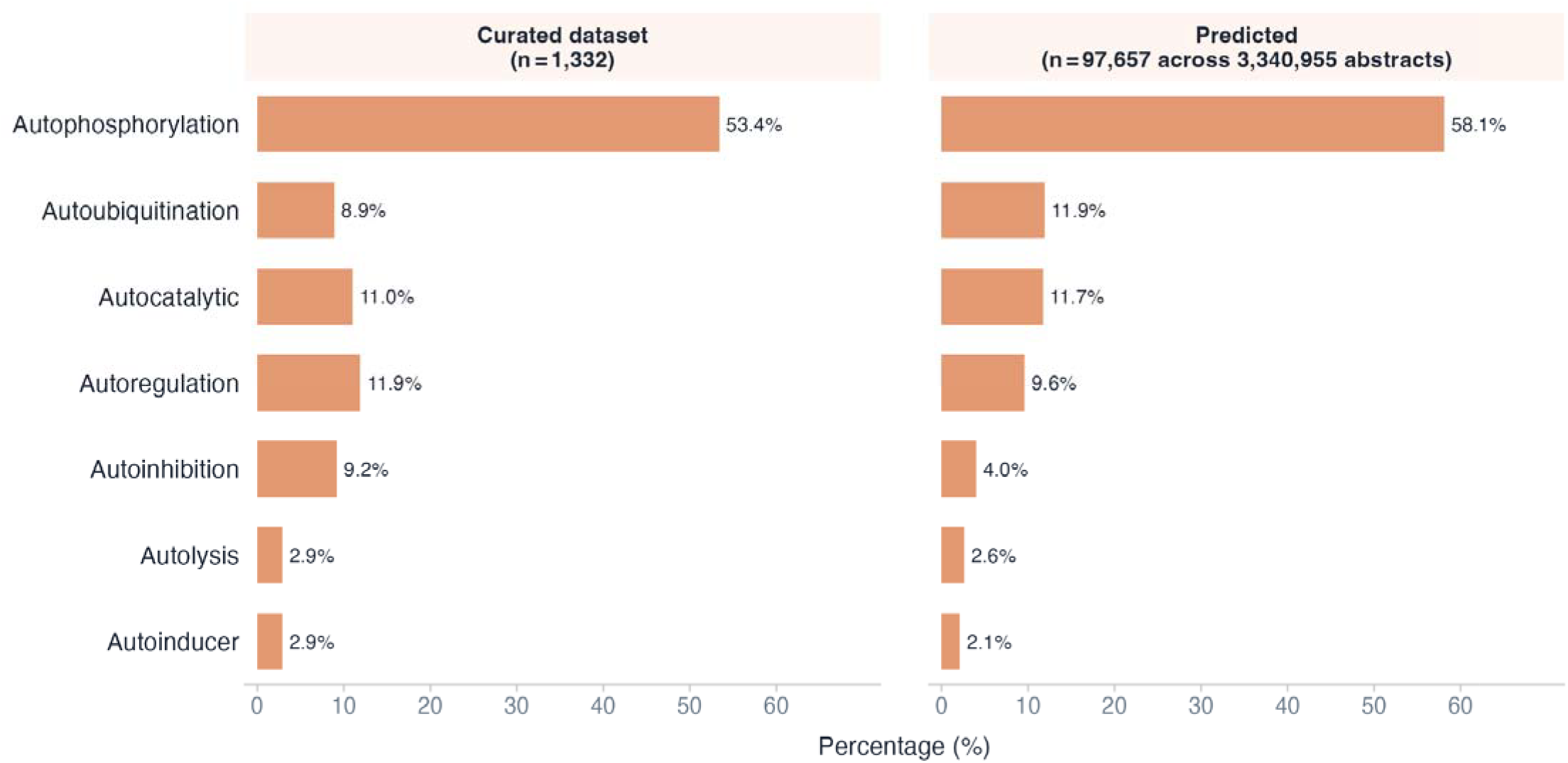
Distribution of predicted autoregulatory mechanism types compared to the curated dataset. Left panel: percentage breakdown of 97,657 mechanism predictions across 3,340,955 unseen PubMed abstracts. Right panel: percentage breakdown of the 1,332 manually curated instances used for model training. Both panels share the same x-axis scale to facilitate direct comparison.

On the Search tab, users can filter by protein identifiers (Protein ID, UniProt AC), organism (OS), PMID, author, journal, data source (Predicted, UniProt, SIGNOR, TRRUST, OmniPath), autoregulatory mechanism, polarity (+/–/±), and publication year range - details for polarity are explained shortly. A free-text box enables case-insensitive keyword search across titles and abstracts. Results are rendered as an interactive table (DT) with sortable columns: AC, UniProt AC, Protein ID, Protein Name, Gene Name, OS, PMID, Title, Abstract, Journal, Authors, Year, Month, Source, Autoregulatory Type, Polarity, Mechanism Probability, and Type Confidence. Probabilities are shown as percentages, and long fields are truncated with a magnifier button for full inspection in a modal. Users can apply confidence cut-offs via the Has Mechanism and Autoregulatory Type controls, then export the filtered results as CSV for downstream review. Each record includes a Polarity indicator derived deterministically from mechanism type: positive (+) for self-amplifying mechanisms (autophosphorylation, autocatalytic, autoinducer), negative (–) for self-limiting mechanisms (autoinhibition, autoubiquitination, autolysis), and context-dependent (±) for autoregulation.

The Statistics tab includes interactive visualizations that display the distribution of autoregulatory types, publication timeline (bubble chart by year), top contributing journals, and data source composition. Histograms of Mechanism Probability and Type Confidence distributions help users select filtering thresholds. Model training performance metrics are also displayed.

The Ontology tab provides the curated vocabulary used by SOORENA. A hierarchical tree presents high-level groupings followed by mechanism pages for the seven classes, each with a prose definition, core relations (is-a, part-of, regulates, has-input, has-output, occurs-in), and key references. From the Search table, each classified record includes a popup that shows the ontology path and the same definition block, keeping model outputs tied to consistent semantics.

The app is versioned through a Patch Notes tab that lists dated changes to data columns, evaluation summaries, UI components, ontology content, and About tab credits contributors and partners. Together, these elements make the system auditable and reproducible: users can trace any row back to its PMID, see the model’s probability for mechanism presence and type, consult standardized definitions, and export subsets for manual validation.

### 3.5 Comparing Source of Autoregulation Data

The SOORENA database integrates autoregulatory data from two complementary sources: literature-derived predictions generated by the two-stage NLP pipeline and entries drawn from four manually curated external databases (UniProt, SIGNOR, TRRUST, and OmniPath). Of the 100,065 total autoregulatory entries, 97,657 (97.6%) originate from SOORENA predictions spanning 3,340,955 PubMed abstracts, while the remaining 2,408 entries (2.4%) are sourced from curated databases (UniProt, n = 1,332; SIGNOR, n = 995; TRRUST, n = 61; OmniPath, n = 20) (**Fig. 8A**). This disparity reflects both the breadth of the literature search and the inherently limited coverage of manually curated resources.

**Figure 8.**
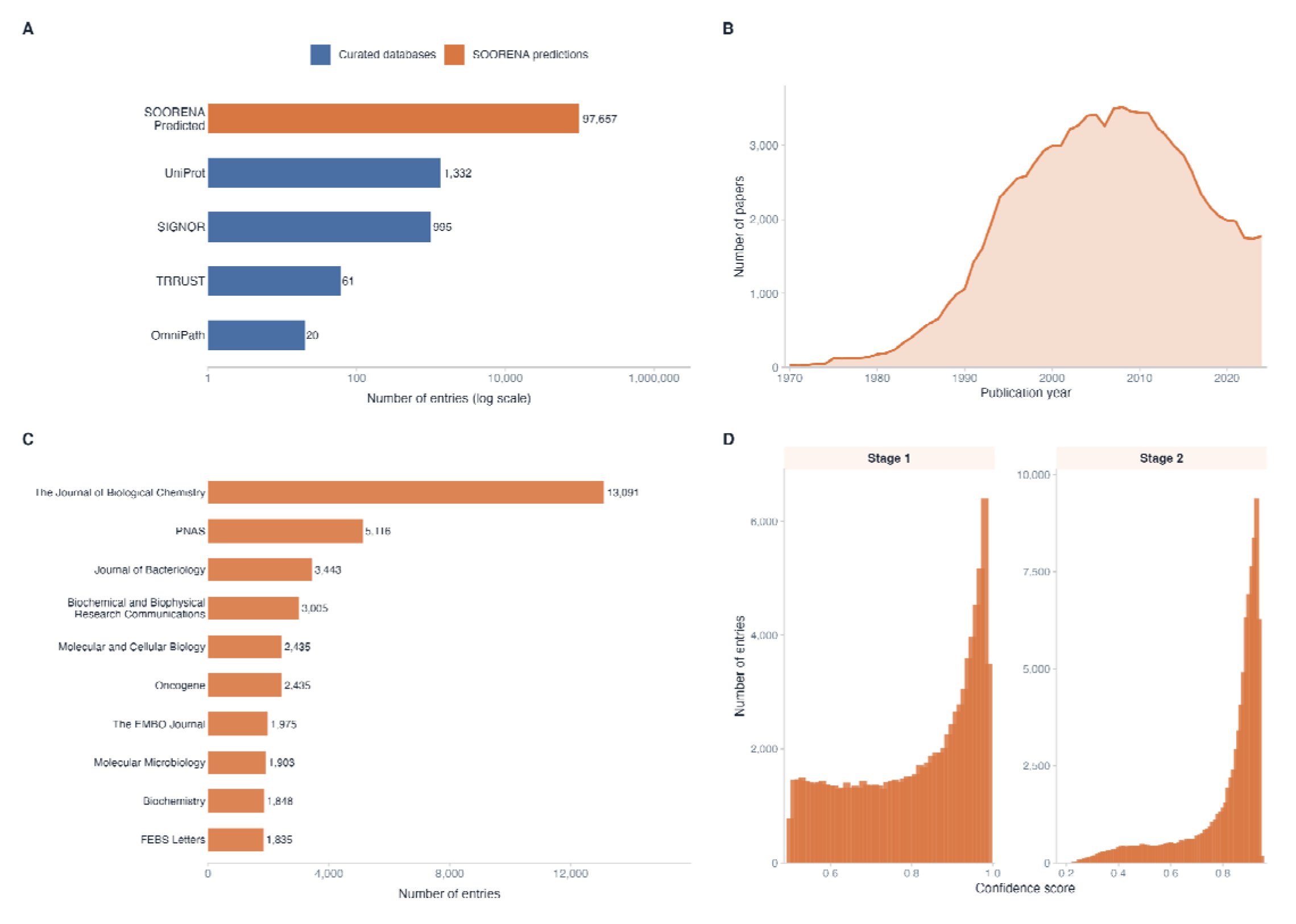
Overview of the SOORENA autoregulation dataset. (A) Number of entries contributed by each data source on a log scale. SOORENA predictions (orange; n = 97,657) account for 97.6% of all entries; curated database contributions (blue) range from 20 (OmniPath) to 1,332 (UniProt). (B) Annual distribution of publications associated with predicted entries (1970–2024), illustrating the growth of autoregulation research over five decades. (C) Top 10 journals by number of associated entries; the Journal of Biological Chemistry (n = 13,091) is the dominant source. (D) Prediction confidence score distributions for Stage 1 (mechanism detection; range 0.50–0.99) and Stage 2 (mechanism classification; range 0.18–0.95). Both distributions are right-skewed, with the majority of entries assigned high-confidence predictions.

Analysis of the publication years associated with predicted entries reveals a sustained growth in autoregulation-related literature from the 1970s onward, peaking in the mid-2000s and plateauing thereafter, with an apparent decline in recent years attributable to publication indexing lag rather than a true reduction in research output (**Fig. 8B**). The Journal of Biological Chemistry contributed the largest number of entries (13,091), followed by PNAS (5,116) and the Journal of Bacteriology (3,443), reflecting the predominantly biochemical and microbiological character of the autoregulation literature (**Fig. 8C**).

Prediction confidence scores for both pipeline stages exhibit right-skewed distributions, indicating that the majority of predictions are made with high certainty (**Fig. 8D**). Stage 1 (mechanism detection) scores are bounded at 0.5 by the binary classification threshold, with a pronounced accumulation of entries above 0.9 and a sharp peak at 0.98. Stage 2 (mechanism classification) scores span a wider range (0.18–0.95), reflecting the greater difficulty of the seven-class task, yet similarly concentrate above 0.8, with a peak at 0.93. Together, these distributions demonstrate that low-confidence predictions represent a minority of the dataset and that the pipeline assigns decisive probabilities across the majority of entries.

## 4. Discussion

This study presents SOORENA, a transformer-based system for automated discovery and mechanistic classification of protein autoregulatory mechanisms in biomedical literature. The two-stage architecture achieved strong performance across all seven mechanism classes, including rare ones, and enabled large-scale deployment across millions of PubMed abstracts. Integrated with experimentally validated annotations from UniProt, SIGNOR, TRRUST, and OmniPath, SOORENA provides the first comprehensive, searchable database of protein self-regulatory mechanisms.

Manual curation remains the gold standard for accuracy but scales poorly and cannot keep pace with the rapid growth of publications [29]. Rule-based text mining can identify explicit mechanistic terms but often fails when descriptions are implicit or context-dependent, which is common for post-translational and feedback processes. SOORENA bridges this gap by combining transformer-based language understanding with domain-specific fine-tuning, achieving expert-comparable interpretation while enabling continuous, automated updates to regulatory knowledge bases.

Transformer models pretrained on biomedical corpora, particularly PubMedBERT, show major advantages in biomedical reasoning over general-domain models such as BERT-base [19,20,30,31]. In preliminary experiments on an earlier formulation of this task, vanilla BERT achieved only 42.6% F1 compared to 94.7% for BioBERT and 98.2% for PubMedBERT, confirming that domain-specific pretraining is essential for biomedical mechanism detection. Our findings reinforce the importance of domain alignment during pretraining, especially when training labeled datasets are small, which is a frequent bottleneck in biomedical NLP.

The staged architecture reflects the structure of the biological question and improves reliability. Stage 1 prioritizes high precision to prevent irrelevant publications from propagating errors downstream, a well-established requirement in hierarchical classification pipelines [23]. Stage 2 uses weighted loss to counter class imbalance, a well-established requirement for biomedical ML tasks where minority classes are often scientifically most important [24]. This formulation addresses the limitations of single-stage classification, where class imbalance and task heterogeneity commonly degrade performance. Additionally, routing only ∼2.5% of publications into Stage 2 reduces computational cost by approximately 94%, enabling scalable whole-corpus inference on standard hardware.

Misclassifications were concentrated among mechanistically adjacent categories. For example, autophosphorylation was occasionally misclassified as autocatalytic activity. Biochemically, these mechanisms are closely related: autophosphorylating kinases are executing a specific form of enzymatic autocatalysis [25,26]. Likewise, confusion among ubiquitin-dependent processes suggests contextual ambiguity in abstracts, where enzymatic roles (E2 vs. E3 ligases) are often unstated or implicitly described.

Crucially, no misclassifications occurred between mechanistically distant biological categories (e.g., autolysis vs. autoinduction), indicating that the model’s representations capture mechanistic semantics rather than relying on lexical co-occurrence patterns. Probability calibration analysis further supports its practical usability: correctly classified predictions displayed higher confidence scores than misclassified ones, allowing confidence-threshold filtering to assist expert curation and quality control review [27].

Reliance on UniProt introduces several known curation biases. Positive examples are skewed toward well-studied proteins and pathways, particularly kinase signaling, thereby inflating the autophosphorylation frequency relative to biological reality. A second key limitation is the use of abstracts rather than full-text articles. Mechanistic assertions often appear in results sections, figures, or supplemental materials [28], meaning that both training labels and predictions are constrained by incomplete contextual information. This likely underestimates true model capability. Handling of multi-mechanism publications remains simplified. Although biologically meaningful co-regulation exists (e.g., autophosphorylation-dependent autoinhibition), only 18 such cases were identified, insufficient for a reliable multi-label model. The model’s reliance on transformer-based semantics inference introduces a potential for latent confounders. For instance, detecting phosphorylation-related language does not guarantee that the event is auto-directed; distinguishing cis-from trans-regulatory contexts remains challenging at the abstract level.

A critical limitation affecting deployed predictions is the system’s inability to identify which specific protein exhibits autoregulatory behavior when multiple proteins are mentioned. The two-stage model predicts whether an abstract describes autoregulation but does not perform subject identification. Gene annotations are extracted post-hoc using PubTator3 from full-text articles, and predictions are paired with all extracted gene mentions. This architecture introduces systematic false positives when abstracts describe heteroregulation (Protein A regulates Protein B): the target protein (B) may be incorrectly flagged as autoregulatory. Pattern-matching analysis identified 3,034 such cases (3.1% of deployed predictions), representing a known error class that requires subject-protein identification in future iterations.

Beyond these limitations, SOORENA provides a scalable framework for expanding biochemical knowledge by identifying regulatory findings that remain uncurated. The integration of 1,076 interactions from pathway databases (SIGNOR, TRRUST, OmniPath) alongside 97,657 literature-derived predictions creates an extensive resource that complements manual curation efforts. High-confidence predictions indicate literature that may contain overlooked experimental evidence, and this helps accelerate the update cycle for resources such as UniProt. The comprehensive mapping of mechanism types across thousands of proteins also enables comparative biological analysis, offering potential insights into how self-regulatory processes have diversified evolutionarily across kinases, proteases, and quorum-sensing systems. Moreover, linking predicted mechanisms with protein domains, structural motifs, and interaction partners may support new hypotheses regarding the molecular features that predispose certain proteins to self-regulate, consistent with prior studies suggesting structural correlates of autoregulation [32,33].

Temporal patterns in autoregulatory discoveries may further reveal research trends and highlight neglected regulatory processes that warrant renewed experimental focus. Finally, integrating SOORENA predictions with structural and functional annotation pipelines could directly support targeted experimental design by identifying proteins most likely to exhibit self-regulatory control and thus prioritizing them for validation.

Future extensions of this work will focus on increasing the granularity, completeness, and biological context of autoregulatory mechanism detection. Incorporating full-text articles instead of abstracts would substantially improve mechanistic recall, as essential experimental details often appear in figure legends, methods, and supplementary materials. Addressing this challenge will likely require long-sequence or hierarchical transformers capable of processing multi-kilobyte documents efficiently [34]. Addressing the subject-protein identification limitation is a priority. Future versions should incorporate entity-aware architectures or sequence tagging approaches to identify which protein in a multi-protein abstract exhibits the autoregulatory behavior, eliminating the current 3.1% cross-regulation error rate introduced by post-hoc gene pairing. Expanding beyond the current single-label formulation to support multi-mechanism prediction is also important since many proteins employ layered regulatory strategies such as autophosphorylation-dependent autoinhibition. This will likely require modified classification architectures and additional annotated examples of co-occurring mechanisms. In parallel, advancing toward knowledge-augmented inference by leveraging protein interaction networks, domain annotations, and curated mechanistic pathways could provide a biologically grounded framework for disambiguating closely related mechanisms. Finally, integrating active learning driven by the model’s well-calibrated uncertainty estimates could make expert validation more efficient, allowing iterative refinement of both the system and the underlying curated knowledge.

## Acknowledgements

We gratefully acknowledge the contributions of Zheng He, Yining Zhou, and Mingyang Zhang in the early prototyping and application development phases of this work. This study was financially supported by the Tampere Institute for Advanced Study and the Jane and Aatos Erkko Foundation [Grant 220031 to M.J.].

## Data and code availability

The script and data are publicly available at (https://github.com/jafarilab/SOORENA).

## Author Contributions

M.J. conceived the research project, and M.J. and P.N. jointly supervised and coordinated its execution. H.A. contributed to the modeling and prepared the initial draft of the manuscript. J.A. contributed to data collection for the training and test sets. All authors participated in writing, editing, and revising the manuscript.

